# Automated speech artefact removal from MEG data utilizing facial gestures and mutual information

**DOI:** 10.1101/2024.09.15.613166

**Authors:** Sara Tuomaala, Salla Autti, Silvia Federica Cotroneo, Pantelis Lioumis, Hanna Renvall, Mia Liljeström

## Abstract

The ability to speak is one of the most crucial human skills, motivating neuroscientific studies of speech production and speech-related neural dynamics. Increased knowledge in this area, allows e.g., for development of rehabilitation protocols for language-related disorders. While our understanding of speech-related neural processes has greatly enhanced owing to non-invasive neuroimaging techniques, the interpretations have been limited by speech artefacts caused by the activation of facial muscles that mask important language-related information. Despite earlier approaches applying independent component analysis (ICA), the artefact removal process continues to be time-consuming, poorly replicable and affected by inconsistencies between different observers, typically requiring manual selection of artefactual components. The artefact component selection criteria have been variable, leading to non-standardized speech artefact removal processes. To address these issues, we propose here a pipeline for automated speech artefact removal from MEG data. We developed an ICA-based speech artefact removal routine by utilizing EMG data measured from facial muscles during a facial gesture task for isolating the speech-induced artefacts. Additionally, we used mutual information (MI) as a similarity measure between the EMG signals and the ICA-decomposed MEG to provide a feasible way to identify the artefactual components. Our approach efficiently and in an automated manner removed speech artefacts from MEG data. The method can be feasibly applied to improve the understanding of speech-related cortical dynamics, while transparently evaluating the removed and preserved MEG activation.

## 1 Introduction

Communication through speech is crucial for humans, and an important part of our wellbeing. Over the last decades, our understanding of the neural dynamics underlying speech production has greatly improved owing to non-invasive studies utilizing electrophysiological techniques such as magnetoencephalography (MEG) and electroencephalography (EEG). Increased knowledge of the cortical functions related to speech and language allows, e.g., for the development of rehabilitation protocols for language-related disorders, and increased precision in presurgical mapping of eloquent speech areas before resection in cases of tumor and drug-resistant epilepsy. However, in basic research and clinical settings utilizing MEG or EEG identifying brain regions vital for speech is hampered by the prominent artefacts produced during speech.

Artefacts during vocal speech originate from various sources, such as the facial muscles and the movements of the head and the jaw (Aristei et al., 2011; Fargier et al., 2018; Ouyang et al., 2016). In addition, the tongue movement produces a prominent artefact, the glossokinetic potential (Vanhatalo et al., 2003). The largest facial muscles activated during speech are the masseter and zygomaticus major, but most facial muscles used for speech are small and largely overlapping in their activity (Stepp, 2012). As the muscle artefacts arise from many independent muscle groups, it is difficult to characterize them and thus challenging to remove them from the MEG and EEG signals (McMenamin et al., 2010; Urigüen & Garcia-Zapirain, 2015). Moreover, muscular activity, as measured with surface electromyography (sEMG), has a wide spectral range from almost 0 Hz to *>* 200 Hz, typically peaking around 100 Hz (Urigüen & Garcia-Zapirain, 2015). The spectral contents of the recorded electrophysiological signals from the brain thus overlap with the EMG activity (Jiang et al., 2019; McMenamin et al., 2010; Urigüen & Garcia-Zapirain, 2015), further complicating the artefact removal.

Picture naming offers a convenient way to study the neural underpinnings of word production (Indefrey & Levelt, 2004; Levelt et al., 1998; Salmelin et al., 1994) since it involves the natural sequence of language production from conceptual processing to word articulation. It is also the most common task used to evaluate speech errors induced by brain stimulation in presurgical evaluation and during awake craniotomy (Corina et al., 2010; Krieg et al., 2017). Even though picture naming and its variations are widely used in both experimental brain research and clinical settings (Kohn & Goodglass, 1985; Krieg et al., 2017; Lioumis et al., 2012; Raffa et al., 2022; Sonoda et al., 2022), the prominent muscle artefacts limit the usage of vocal speech tasks during MEG and EEG recordings. Many attempts have been made to overcome this problem by modifying the task to avoid oral responses. For example, delayed (Ala-Salomäki et al., 2021; Jescheniak et al., 2002), silent (Eulitz et al., 2000; Liljeström et al., 2009), whispering (Eulitz et al., 2000), and manual responses (Schmitt et al., 2000) have been used. In addition, some studies have limited the data analysis to the time window before the onset of the speech response, when muscular artefacts do not yet contaminate the measured brain activation (Habets et al., 2008; Liljeström et al., 2015). Yet, despite the prominent muscular artefacts, using a vocal naming response has remarkable experimental advantages as it tracks the full word production process and ensures that one can assess whether the subject follows the given test instructions and names the pictures correctly. The need to remove the muscular contamination from the brain signals in vocal naming thus prevails.

Different methods for speech artefact removal have been proposed, such as removing the high-frequency muscular signal from the recordings by a low-pass or band-pass filtering (Ganushchak & Schiller, 2008; Laganaro & Perret, 2011), or applying blind source separation by canonical correlation analysis (Vos et al., 2010) to identify and remove the artefactual components. Additionally, algorithms utilizing independent component analysis (ICA), with different component selection criteria have been previously suggested. Such criteria have included, e.g., statistical characteristics like kurtosis (Barbati et al., 2004; Porcaro et al., 2015), correlation of the components with the concurrent EMG signals (Alexandrou et al., 2017; Porcaro et al., 2015; Vos et al., 2010) or their power spectrum densities (PSDs) (Barbati et al., 2004; Porcaro et al., 2015), visual inspection of the component topography (Alexandrou et al., 2017; Hipp et al., 2011; Kujala et al., 2013; Porcaro et al., 2015), component localization at the source level (Alexandrou et al., 2017; Henderson et al., 2013; Porcaro et al., 2015), and the relative amount of gamma (∼ 40 Hz)/high-gamma (*>* 80 Hz) activation in their spectral power (Alexandrou et al., 2017; Hipp et al., 2011). Despite the proposed methods, muscle artefact removal from speech data remains challenging and time-consuming. With manual component selection criteria the artefact removal process may be poorly replicable and hampered by problems in inter-observer reliability (Gross et al., 2013; Muthukumaraswamy, 2013). Conversely, the more automatized selection criteria tend to vary between studies leading to non-standardised speech artefact removal processes. Evidently, there is a need for a reproducible and automatized speech artefact removal pipeline, with rather simple component selection criteria.

This study aims to improve the artefact removal process and proposes a replicable pipeline for automated speech artefact removal from MEG data. We designed a facial gesture task to be performed before the actual naming task for collecting examples of the most prominent facial muscular artefacts. The facial gesture task contains 10 movements that use the muscles essential to speech; muscle activity is measured over four facial muscles with EMG. Component selection is automated using mutual information (MI) between the EMG signals and the ICA-decomposed MEG signals. The use of MI as a similarity measure is motivated by its ability to find non-linear dependencies between two signals and its better resilience to outliers compared to correlation (Correa & Lindstrom, 2013). Similar approaches have shown promise for extracting deep brain stimulation (DBS) artefacts from MEG data (Abbasi et al., 2016) and cardioballistic artefacts from EEG signals (Abbasi et al., 2015; Liu et al., 2012). As a similarity measure, MI requires setting a threshold to define when two signals are considered similar enough. To solve this issue, we employed k-means clustering. We compared two different approaches: Mutual Information (MI) with clustering (C), denoted as MIC-ICA, and manual classifying of the artefactual components (Manual-ICA).

We evaluated whether the addition of a facial gesture task and the use of MI as a similarity measure can improve the artefact removal process in healthy subjects performing a picture naming task. The results were assessed from two perspectives: firstly, by evaluating how well the artefact removal process preserves true brain activation, and secondly, by measuring the success of the automated artefact removal process. Preservation of brain activation was evaluated by comparing cleaned data from a vocal naming task to that from a silent naming one, whereas the evaluation of the automation process was based on a comparison between the Manual-ICA and the proposed MIC-ICA approach. Our results suggest that using the facial gesture task and simultaneous EMG measurements enables effective automation of the speech artefact reduction process in language-related MEG tasks with vocal speech responses, such as picture naming.

## 2 Materials and Methods

Figure 1 illustrates the overall study paradigm and the analysis workflow. First, all participants took part in a facial gesture paradigm, followed by a standard picture naming paradigm, which included vocal and silent naming tasks and a picture observation task. Hereafter, the picture naming paradigm refers to all these three subexperiments together (vocal naming, silent naming, and observation), while vocal and silent naming tasks refer specifically to the individual tasks. Electrocardiogram (ECG) and electrooculogram (EOG) were recorded simultaneously with the EMG and MEG data. The data was first pre-processed (see below for details) such that prominent external noise originating outside the helmet, eye blinks, and heart artefacts were removed, while the speech artefact remained. The differences between the three speech artefact rejection approaches were then analyzed. These approaches included two automated rejection methods that utilized mutual information (MI) and clustering (C) (MIC-ICA) to select the artefact components. In MIC-ICA-N, the ICA decomposition was applied to the naming paradigm data alone, while in MIC-ICA-GN the ICA decomposition was applied to both the gesture and naming paradigm data. In the MIC-ICA approaches, component selection used mutual information (MI) between the measured facial EMGs and independent components as the metric for k-means clustering to automatically classify the components to be removed. For comparison, as a reference method, the artefactual components were classified manually based on visual inspection of their time series and topographies (Manual-ICA).

**Figure 1.**
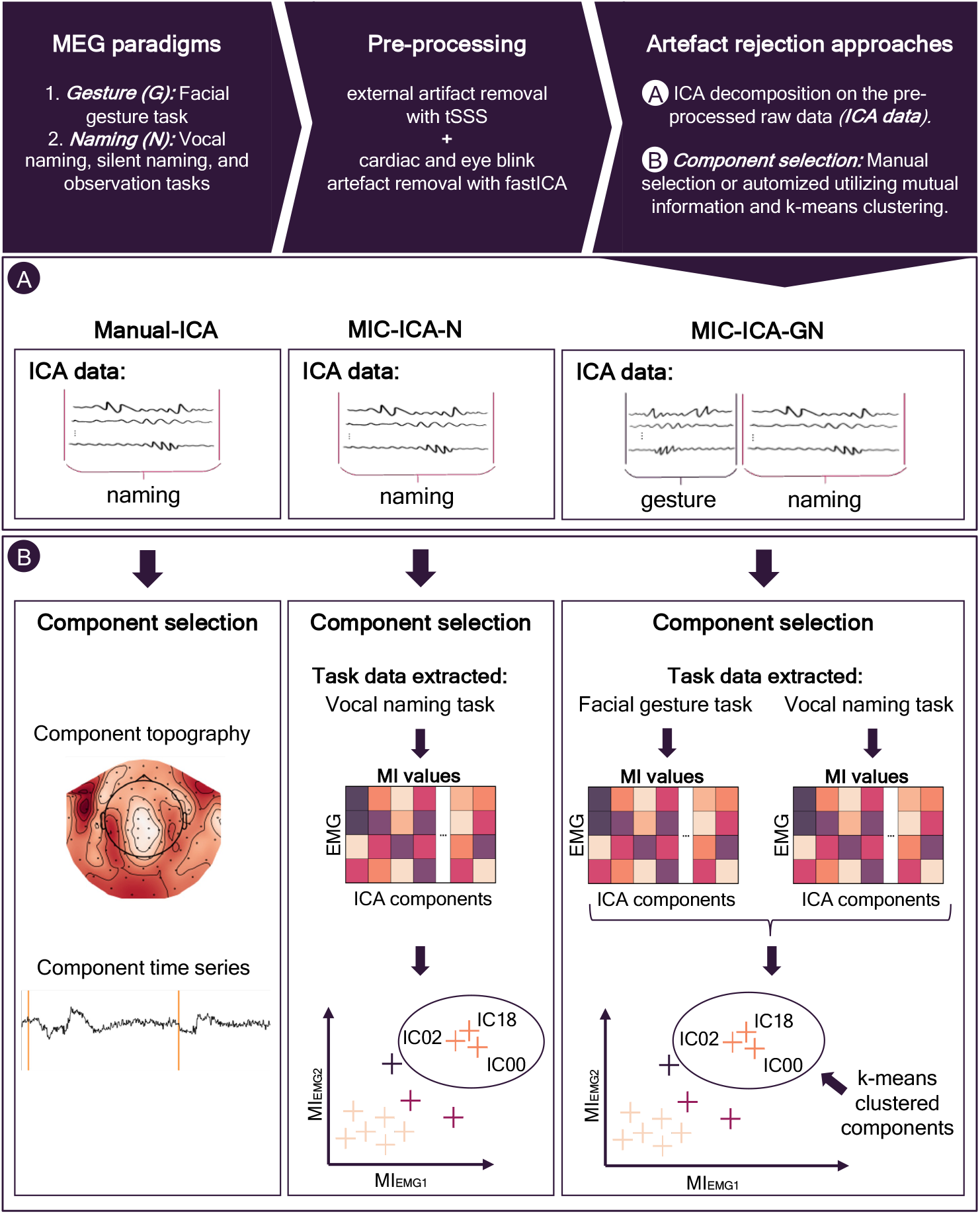
The study pipeline. The study consisted of a facial gesture paradigm (G) and a standard picture naming paradigm (N), consisting of vocal naming, silent naming and observation tasks. The MEG data was accompanied by EMG, EOG and ECG recordings. The measured data was pre-processed using tSSS and FastICA to remove external noise, eye blinks and heart artefacts. To remove speech artefacts, FastICA was applied again, now to the pre-processed data: Manual-ICA and MIC-ICA-N used the data from the naming paradigm, and MIC-ICA-G additionally included the data from the gesture paradigm. In the Manual-ICA approach, the artefact ICA components were selected manually. In the MIC-ICA methods, mutual information values between the EMG signals and ICA-decomposed MEG signals were calculated. Components sharing high MI values were then identified with k-means clustering and subsequently removed.

### 2.1 Subjects

12 healthy volunteers [mean +-STD age 25 +-3 years, range 21-29, 6 females] participated in the study. All subjects were right-handed, without neurological or psychiatric disorders and had Finnish as their native language. The measurement protocol was approved by the Helsinki University Hospital (HUS) Regional Committee on Medical Research Ethics, and all participants signed a written consent form to take part in the study.

### 2.2 Experimental design

#### 2.2.1 Facial gesture task

Before the picture naming paradigm, the participants performed a facial gesture task designed to aid in the automated identification of artefacts. Ten facial movements were performed, each designed to preferentially activate a specific facial muscle. These movements covered gestures that are known to produce strong but non-isolated muscular activations (O’Dwyer et al., 1981; Stepp, 2012, see Table 1).

**Table 1:**
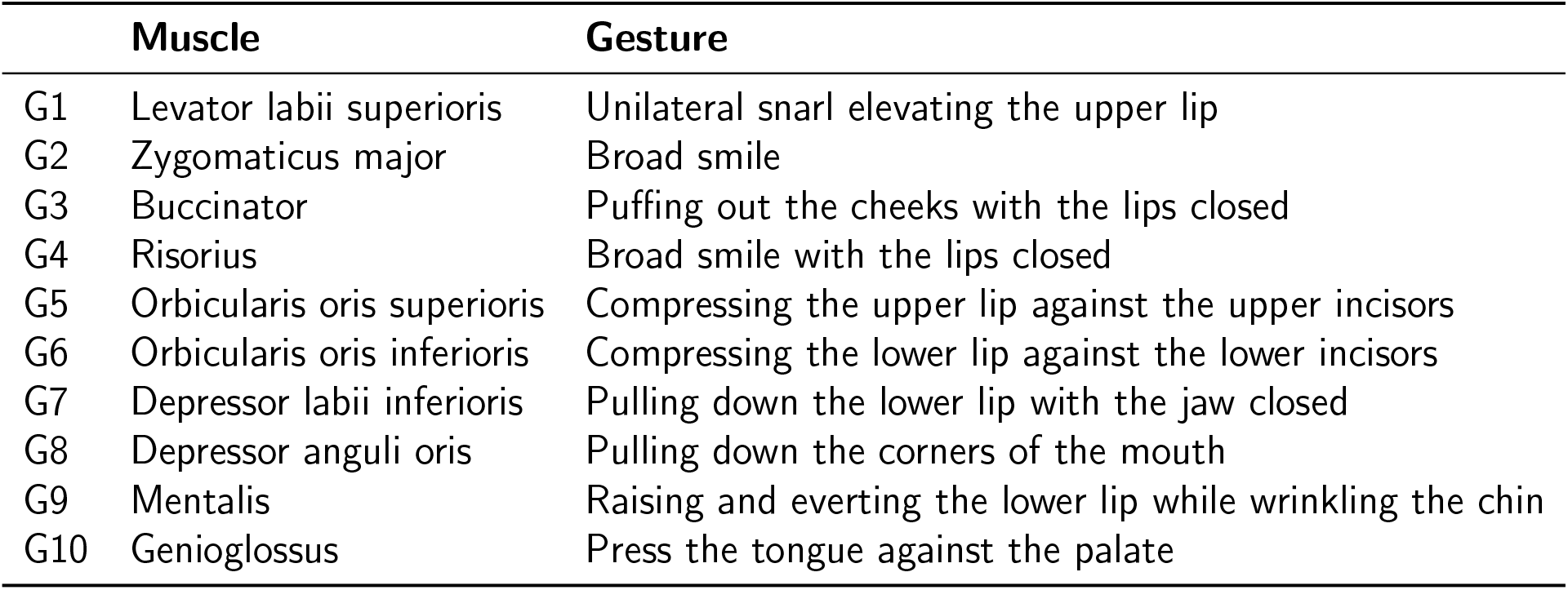
The major muscles activated during speech and the gestures in which they are preferentially used (O’Dwyer et al., 1981).

A block diagram of the facial gesture task is presented in Figure 2. The overall task (Level 1) consisted of five sets of equal length (160 s). Within each set (Level 2), each of the 10 gestures, listed in Table 1, was performed in a randomized order. Each of the gestures within a set was repeated 5 times (Level 3). The subjects were prompted with an audio stimulus to perform the gesture.

**Figure 2.**
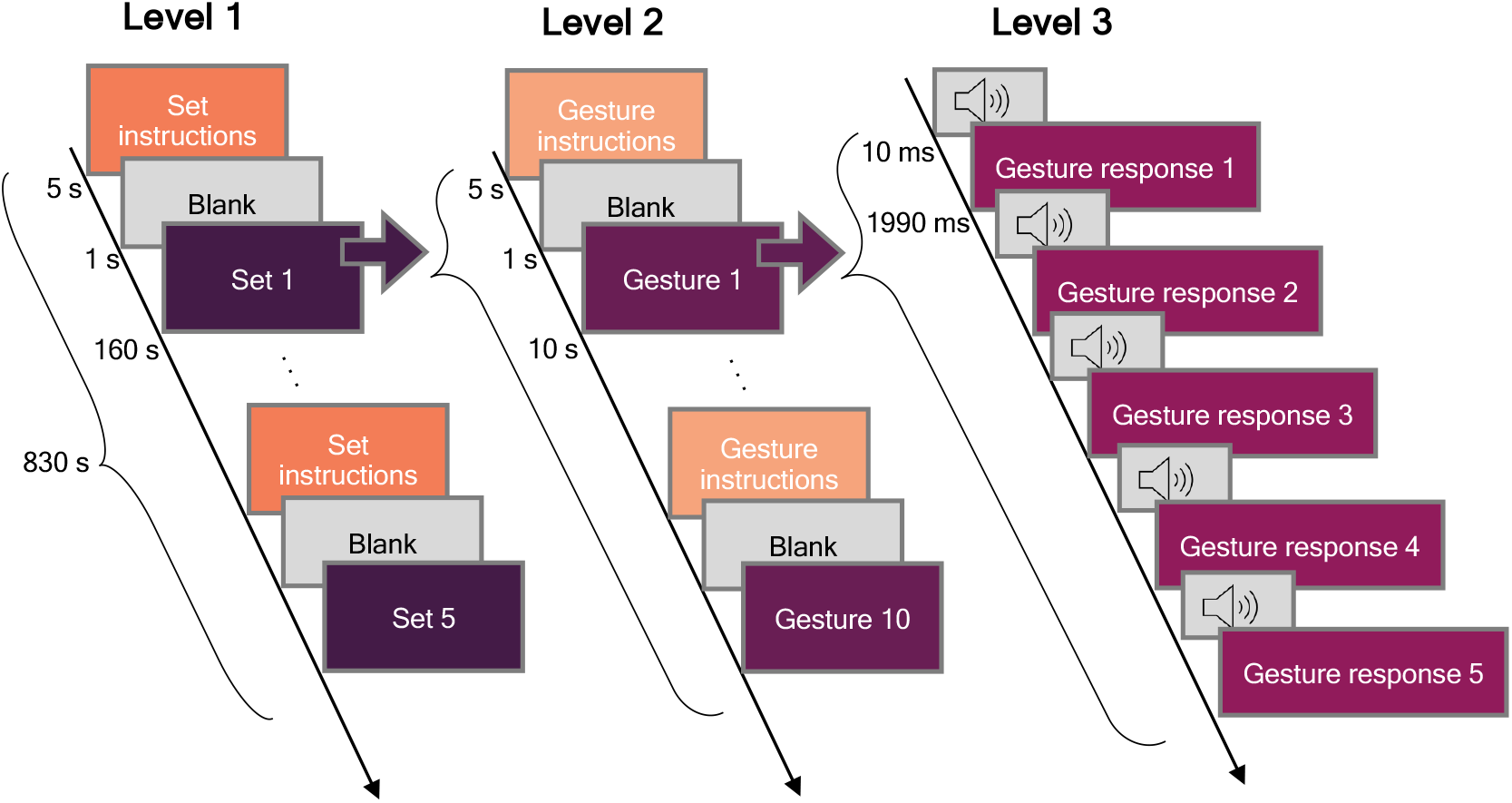
Gesture paradigm structure. The structure of the facial gesture task consists of three levels, depicted as columns in the figure. There were a total of 5 sets in each measurement (Level 1): Within each set, each gesture was presented in a random order (Level 2), and the same gesture was repeated 5 times in a row (Level 3).

#### 2.2.2 Picture naming paradigm

All subjects performed a picture naming paradigm after the facial gesture task using normed colour images of objects (Brodeur et al., 2010, 2014). The structure of the picture naming paradigm is presented in Figure 3. The overall paradigm consisted of three experimental runs (Level 1 presents one run). Within each run, the vocal naming, silent naming, and observation tasks were repeated twice each. The order of these six task blocks was pseudorandomized using three different predetermined configurations. During vocal naming, subjects were instructed to say clearly and fluently aloud the name of the object in the picture presented to her/him. In the silent naming condition, the naming was performed in the subject’s mind without vocal utterances. In the observation condition, no response to the presented picture was required. Three of the subjects did not perform the observation task. During each task, 18-19 pictures were presented (Level 2) and named according to the task. In the vocal and silent naming tasks, each image was shown for 500 ms, after which there was a 2500 ms to respond, followed by a fixation cross for 1000 ms before the next stimulus (Level 3). In the observation task, each image was shown for 500 ms, and the observation period that followed the figure lasted only 1500 ms. In the three runs in total, 110 images were shown during each task.

**Figure 3.**
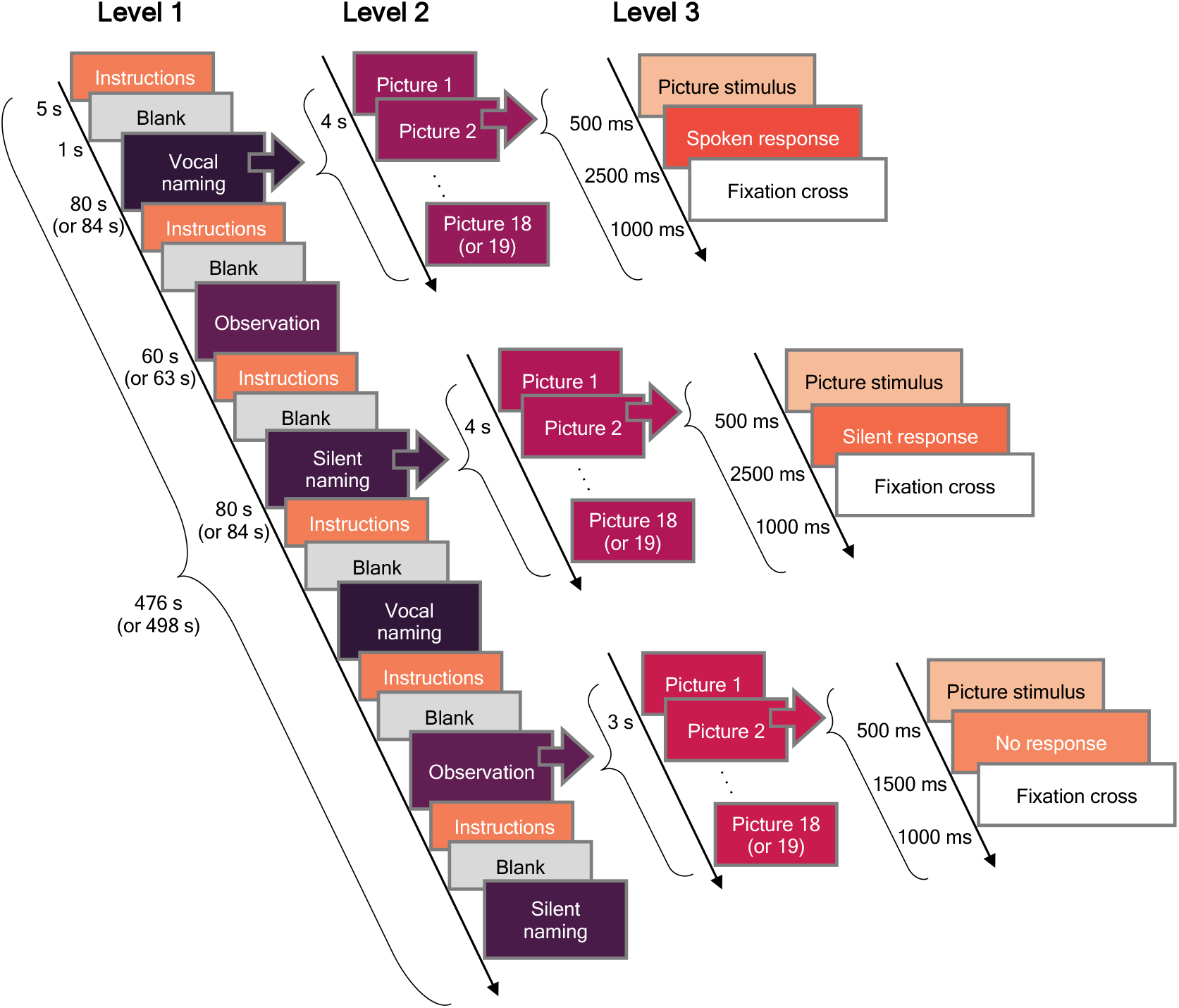
Picture naming paradigm. The paradigm consisted of three runs, i.e., Level 1 was repeated 3 times. During each task (vocal naming, silent naming, and observation), 18-19 pictures were shown (Level 2). The number of the pictures depended on the run. Picture stimulus and response durations are shown at Level 3.

### 2.3 Recordings

The MEG recordings were conducted in the BioMag Laboratory in HUS Diagnostic Center using a 306-channel whole-head MEG device (Elektra Neuromag TRIUX, MEGIN Ltd, Helsinki, Finland) placed in a magnetically shielded room (Euroshield, Eura, Finland). The device contains 102 three-channel units, each consisting of one magnetometer and two planar gradiometers. The sampling frequency was 1000 Hz with a bandpass of 0.03-330 Hz. Five head position indicator (HPI) coils were attached to the face and scalp, and their locations were digitized with a 3-D digitizer pen (Fastrak, Polhemus, US). The 3-D digitizer was also used to digitize the subject’s head shape and three anatomical landmarks (the nasion and the right and the left preauricular points). To track the subject’s head position, the HPI coil positions were measured continuously during the whole MEG measurement by feeding a small current into them. Structural MRI images were acquired at either Aalto Magnetic Imaging Centre (Aalto University, Espoo) or at HUS with a 3T Siemens Skyra MRI scanner.

ECG and EOG were recorded alongside the EMG and MEG data. Two EOG sensor pairs for recording the horizontal and vertical eye movements were attached according to Figure 4.

**Figure 4.**
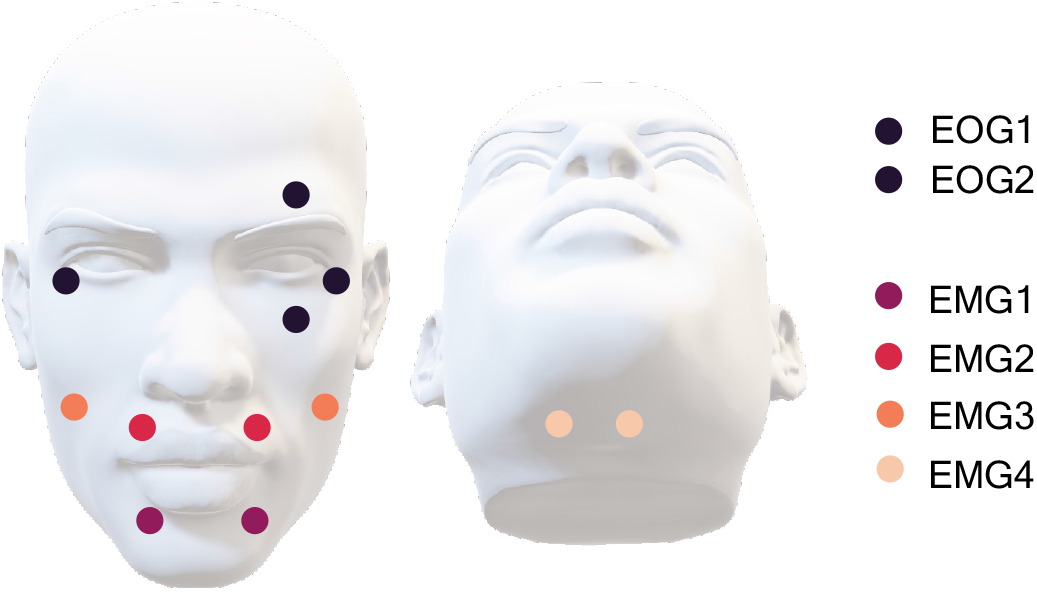
Placements of the EOG and EMG sensors. EOG1 and EOG2 represent the placements of the electrooculogram sensors that measured the horizontal and vertical eye movements. The facial muscle activity was measured with sensors EMG1-EMG4.

Muscle activation was recorded with four surface EMG sensor pairs. The positions of the sensors are visualized in Figure 4. Two EMG sensor pairs were placed above the orbicularis oris superioris (EMG1) and inferioris muscles (EMG2), respectively, similarly to (Abbasi et al., 2021). The third EMG sensor pair was placed above the zygomaticus muscle, and the fourth pair under the chin, aiming to measure the muscle activity close to the rim of the MEG sensor array and activity arising from the tongue, respectively.

### 2.4 Data pre-processing

First, the temporally extended signal space separation (tSSS) algorithm (Taulu & Kajola, 2005) was applied to the data. The method corrects head movements during the MEG measurement and removes noise emerging from sources outside the sensor array. Data from all subjects were then transformed to a common head position. Subsequently, the FastICA algorithm (Hyvärinen, 1999) implemented in MNE Python (Gramfort et al., 2013) was used to remove artefacts originating from the heart and eye movements. Only gradiometer data was used in the analysis.

### 2.5 Speech artefact removal

We compared three different methods for removing the speech artefact. The manual approach (Manual-ICA) provided the reference for the two automated speech artefact removal methods, MIC-ICA-N and MIC-ICA-GN. The three methods differed in terms of which data was used for the ICA decomposition, and how the artefactual independent components (ICs) were selected, as summarized in Figure 1.

#### 2.5.1 ICA decomposition

ICA (Hyvärinen, 1999, 2013; Hyvärinen & Oja, 2000) is widely used as an artefact removal method, also for speech artefacts (Abbasi et al., 2021; Alexandrou et al., 2017; Bourguignon et al., 2020; McMenamin et al., 2010; Porcaro et al., 2015). In this work the ICA decomposition for the speech artefact removal was conducted using the FastICA algorithm in MNE Python (Gramfort et al., 2013; Pedregosa et al., 2011). The algorithm pre-whitens the MEG data with principal component analysis (PCA). To avoid information loss, we required 99.99% of the variance in the data to be explained by the PCA decomposition, resulting in approximately 68 *±* 3 components per participant.

#### 2.5.2 Manual-ICA for a reference approach

In the reference approach, the ICA model was fit using the data from the picture naming paradigm (including the vocal naming, silent naming and observation tasks). The criteria to exclude the artefact components were: 1) visual inspection of the component time series, and 2) visual inspection of the component topography. See Figure 5 for an example of a typical speech artefact time series and topography.

**Figure 5.**
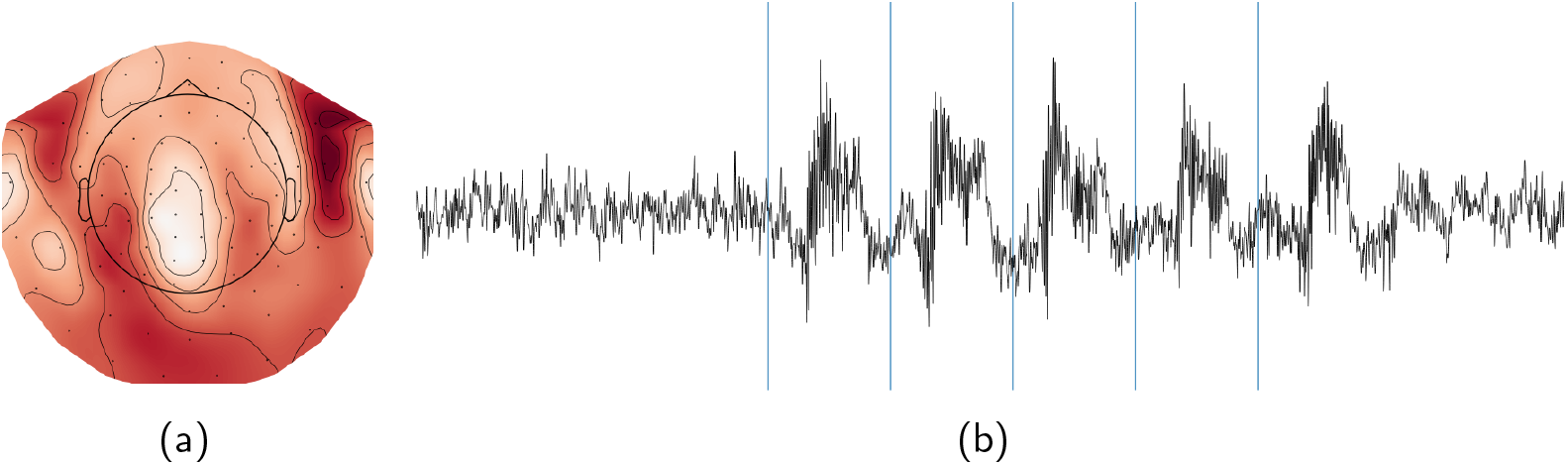
Speech artefact ICA component topography and the corresponding time series. Figure 5a presents the typical topography of a ICA component related to a muscle artefact, and Figure 5b presents the time series of the component.

#### 2.5.3 MIC-ICA approaches

The MIC-ICA approaches aimed to automate the selection of the artefactual ICA components. The ICA decomposition was based on the FastICA algorithm, similar to the manual approach. The two MIC-ICA approaches utilized different data in their ICA decompositions: MIC-ICA-GN utilized both the gesture and naming paradigm data and MIC-ICA-N only the naming paradigm data. The ICA decomposition was made for the entire raw data time series, thus including the vocal naming, silent naming and observation tasks. Hereafter the observation task data was not used in the analysis.

The automated component selection was based on the mutual information values between epochs of the PCA-decomposed EMG signals and the ICA-decomposed MEG sources. The MI values were estimated separately for the ICs of the facial gesture and vocal naming tasks and their respective EMG signals. Therefore, in the case of MIC-ICA-GN, this resulted in 8 (EMG PCA components) times ∼ 68 (ICA components) MI values. The MIC-ICA-N approach, which utilized only the data from vocal naming task, resulted in 4 (EMG PCA components) times ∼ 68 (ICA components) MI values. For calculating MI, we used the Ennemi package implemented in Python (Laarne et al., 2021). Here MI is estimated starting from its definition

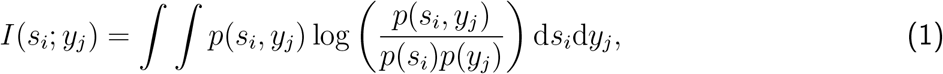

where *p*(*s*_*i*_, *y*_*j*_) is the joint probability distribution between the independent component *s*_*i*_ and the PCA-decomposed EMG component *y*_*j*_ and their marginal distributions are *p*(*s*_*i*_) and *p*(*y*_*j*_), respectively. Mutual information takes values over [0, ∞], and can be normalized to values ranging over [0, 1] in different ways. Here we implemented such normalization by introducing the MI correlation coefficient (Laarne et al., 2022):

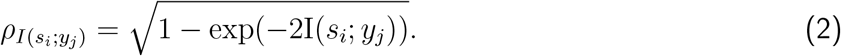

Finally, the components were clustered using the k-means clustering implemented in the Python Scik-itlearn clustering module (Pedregosa et al., 2011) to identify artefactual components. We tested the clustering approach by varying the number of clusters from 2 to 20, finally selecting a cluster number of 5. The number of clusters was selected by visually inspecting the artefact and non-artefact clusters, confirming the decision with the elbow method, which calculates the sum of the squared distances between data points and their closest cluster centre. The cluster number was selected such that adding more clusters no longer yielded significant improvements of the artefact identification. The independent components were ordered according to how high the MI value was for all the EMG sensors (sum of the MI values per IC). All ICs that belonged to the cluster with the highest MI sum were then selected for removal.

### 2.6 Source estimation

Source estimation was performed using MNE Python. Co-registration was performed to match the individual MRI volume and MEG sensor locations. A boundary element model (BEM) was constructed to model the conductivity profile of the head, and a source space of 2562 sources per hemisphere (ico4) was set up to define the possible current source locations. The results from the coregistration, the BEM model and the source space were used to create a forward model. A noise covariance matrix was calculated using the pre-processed epoch baselines from the silent-naming task (110 trials per participant). A source estimate was obtained using noise-normalized cortically constrained minimum norm estimates (dSPM) (loose constraint = 0.3, depth weighting = 0.8). For group-level analysis, the source estimates of each subject were morphed to the Freesurfer ‘fsaverage’ head model.

### 2.7 Evaluation of artefact removal methods

For each of the three artefact removal approaches, the vocal naming task data was reconstructed such that the artefact components obtained by each of the three methods were projected out from the data. Hereafter, we refer to this as the artefact-free data. The silent naming data was analyzed in a similar manner, by projecting out the artefact components obtained by each of the three methods, referred to as the artefact-free silent naming data. This procedure was conducted even though the artefact components were not assumed to show substantial activation during the silent naming. In addition, we used the original vocal naming and silent naming data without any component exclusion in the evaluation.

How well different methods preserved the brain activation was assessed by comparing the original and artefact-free evoked responses in the vocal and silent naming tasks. We assumed that there should be almost no speech-induced artefact present during the silent naming task, and thus the artefact-free and original silent naming evoked responses should not differ considerably. Furthermore, we assumed that the artefact-free and original vocal naming evoked responses would deviate from each other mainly after the speech onset. Accordingly, the less signal is (erroneously) removed before the onset of the speech, the better the brain activation is assumed to be preserved.

#### 2.7.1 Statistical testing

We tested, both at the sensor and source-levels, whether the artefact-free and original vocal and silent naming data statistically differed from each other. At the sensor-level, we tested all sensors independently in the time window of 0-1500 ms. We used non-parametric cluster-level statistical permutation test (Maris & Oostenveld, 2007; Sassenhagen & Draschkow, 2019) from MNE Python version 1.3.0 (Gramfort et al., 2013) with 1000 permutations to control for multiple comparisons; the statistical significance level was set to the p-value of 0.001. The permuted epochs were individually scaled between 0 and 1, by dividing by the (individual) maximum epoch value over all channels.

The statistical significance at the source level was tested with a non-parametric cluster-level permutation test to control for multiple comparisons (Maris & Oostenveld, 2007; Sassenhagen & Draschkow, 2019) with 1000 permutations; the statistical significance level was set to a p-value of 0.05. We tested the difference between the original and cleaned vocal and silent naming task responses from the morphed source estimates, which were normalized by dividing by the maximum (individual) whole-head value. Additionally, we tested if the cleaned vocal naming responses obtained after the different artefact removal methods differed from each other.

#### 2.7.2 Measuring the removed activation

Finally, to measure how much activation was removed in the artefact removal process, we calculated the root mean square deviation (RMSD) between the original and artefact-free evoked vocal naming data at the sensor-level, independently for each speech artefact removal method. First, epochs were extracted from the vocal naming data and averaged to estimate the evoked responses. Evoked responses were windowed 0 - 1500 ms with respect to the image onset, aiming to include most of the spoken response into the averaged signal. Each MEG gradiometer channel pair were then combined using root mean square, after which the evoked responses were normalized by dividing them by the individual whole head maximum. The *RMSD*_*ch*_ were calculated for each channel pair *ch* independently as follows:

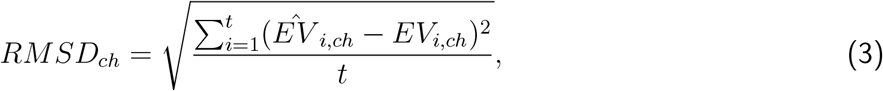

where the original evoked response is marked as 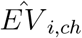, and the evoked response after artefact removal is marked as *EV*_*i,ch*_. Here *i* is the index of the datapoint spanning from 1 to *t* datapoints in the evoked response. A lower *RMSD*_*ch*_ value indicates greater similarity between the original and artefact-free evoked responses.

## 3 Results

### 3.1 Cortical activation during vocal and silent naming

The averaged source-level activations for the uncleaned data and for the data obtained by the different cleaning methods are depicted in Figures 6a (vocal naming) and 6b (silent naming). Overall, the cleaned responses in the vocal naming task are largely similar regardless of the artefact removal method, but differ considerably from the original non-cleaned responses. The most notable differences between the original and cleaned source estimates were observed bilaterally in the anterior temporal and frontal cortices. Importantly, the source-level results for the silent naming data after cleaning were highly similar to the original ones, suggesting that the artefact removal retains the relevant language-related activations. Statistically significant differences between the cleaned and original source estimates are presented in Figure 7. As Figure 7a shows, the brain areas that differ statistically vary slightly between the artefact removal methods: for the MIC-ICA-GN, there is no statistically significant difference in the left frontal brain areas in contrast to MIC-ICA-N and Manual-ICA. However, when testing the statistical differences between the cleaned vocal responses of different methods, none of the tested pairs (MIC-ICA-GN vs. MIC-ICA-N, MIC-ICA-GN vs. Manual-ICA, MIC-ICA-N vs. Manual-ICA) showed statistically significant differences (*p <* 0.05). No statistical differences (*p <* 0.05) were observed between the cleaned and original silent naming data (see Figure 7b).

**Figure 6.**
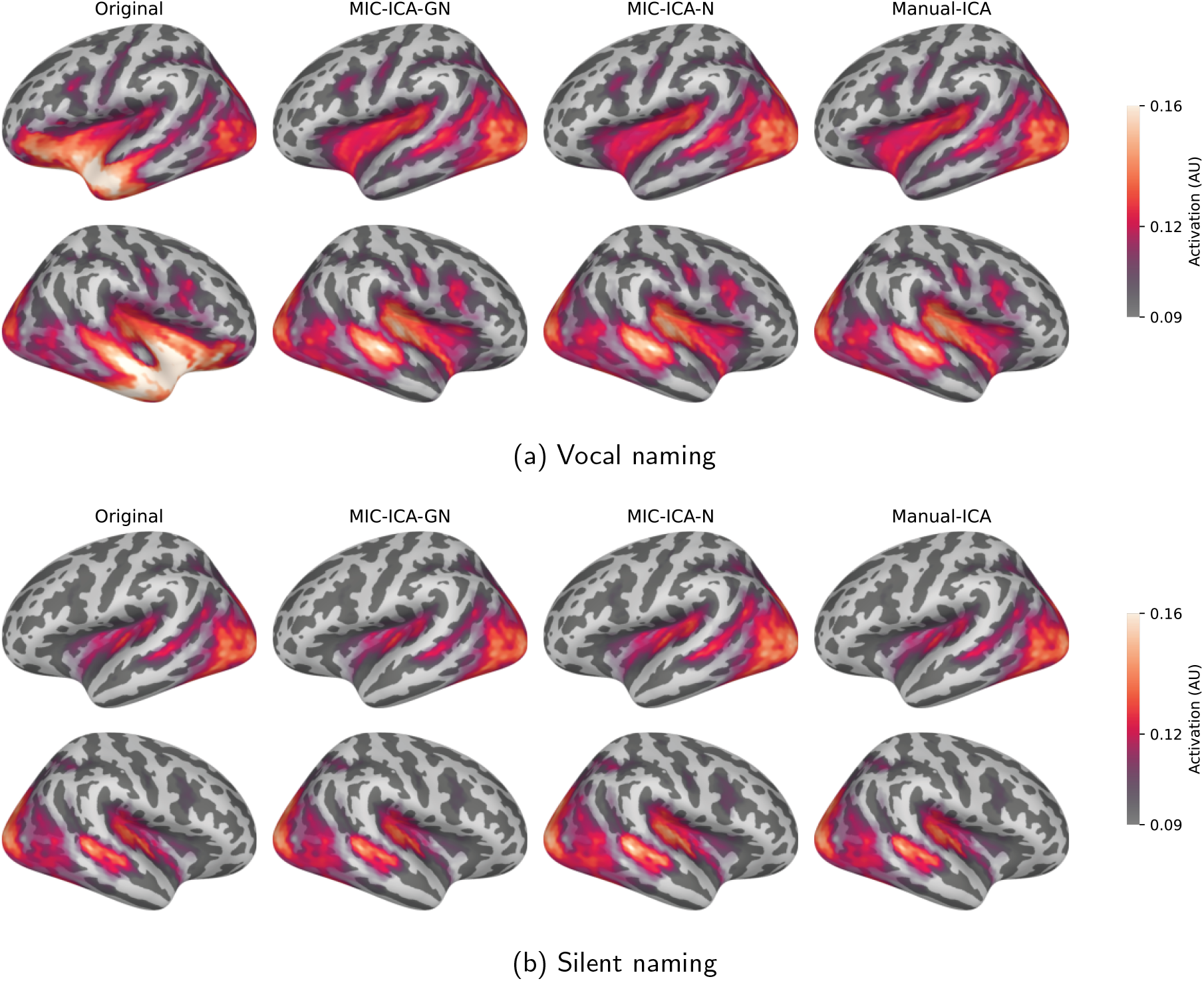
Source-level results in the vocal and silent naming tasks. Source-level results in the vocal (6a) and silent naming (6b) tasks, averaged over all subjects over the time interval between 0 and 1500 ms. The leftmost column shows the original brain responses and the three other columns the cleaned activity after the three different artefact removal methods.

**Figure 7.**
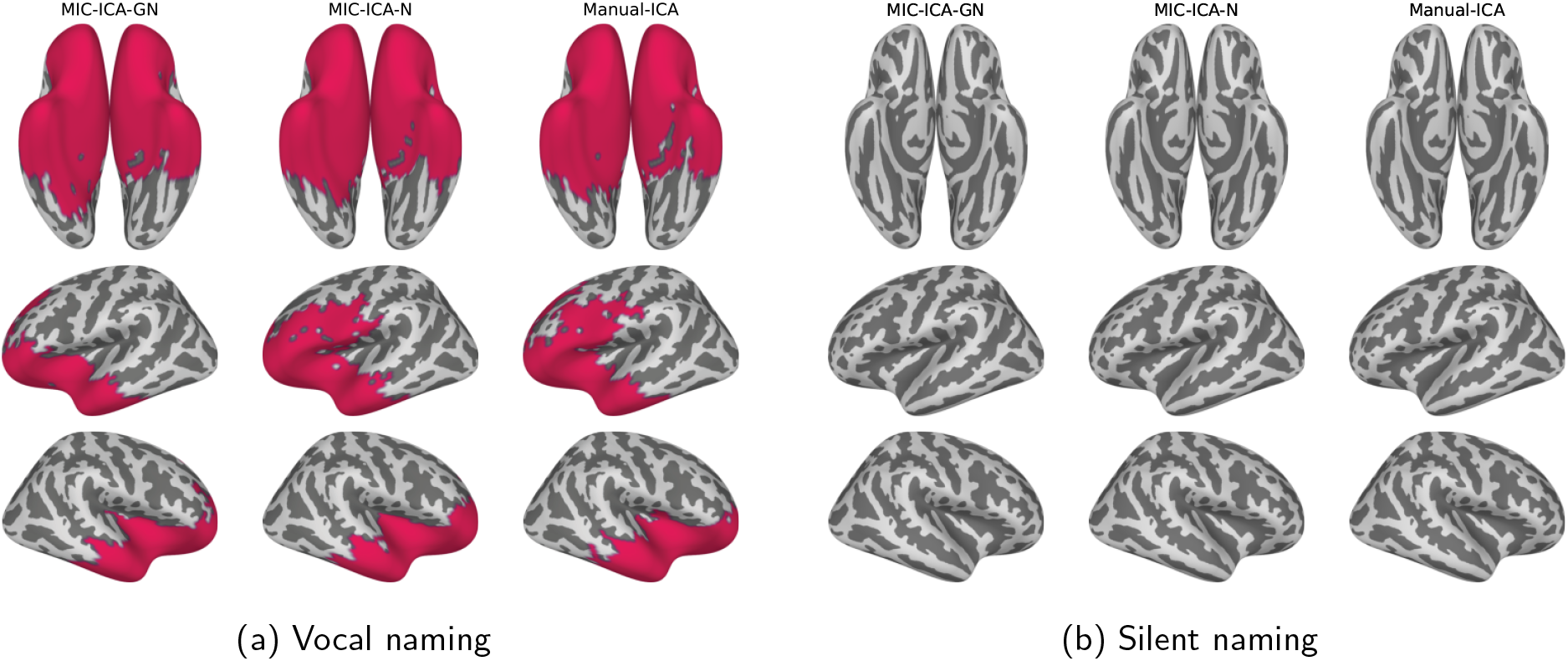
Original vs. cleaned data in the vocal and silent naming tasks. The coloured areas visualize the source areas that differed statistically (*p <* 0.05) between the cleaned and original data. All the areas with statistically significant differences between 0 and 1500 ms are highlighted. No statistically significant differences were observed between the cleaned and original data in the silent naming task.

Despite similarities at the group level, the differences between the artefact removal approaches became apparent at the individual level. Changing the number of clusters *k* was reflected in how many components were classified as artefactual, and therefore assigned to be removed. The average and median number of removed components are presented in Table 2, together with a range of how many components were removed across all subjects. Typically, when a high number of components (*>* 4) were assigned to be removed, visual inspection showed that all of these did not resemble typical speech artefact components. Therefore, the number of clusters was selected such that it would not classify too many components as artefactual and thus set to *k* = 5. Based on the median, average and min.-max. range, it seems that the MC-ICA-GN is slightly more robust, since already with *k* = 3 clusters the MIC-ICA-GN removes a maximum of 4 components across all subjects and altogether the MIC-ICA-GN removes, on average, fewer components.

**Table 2:**
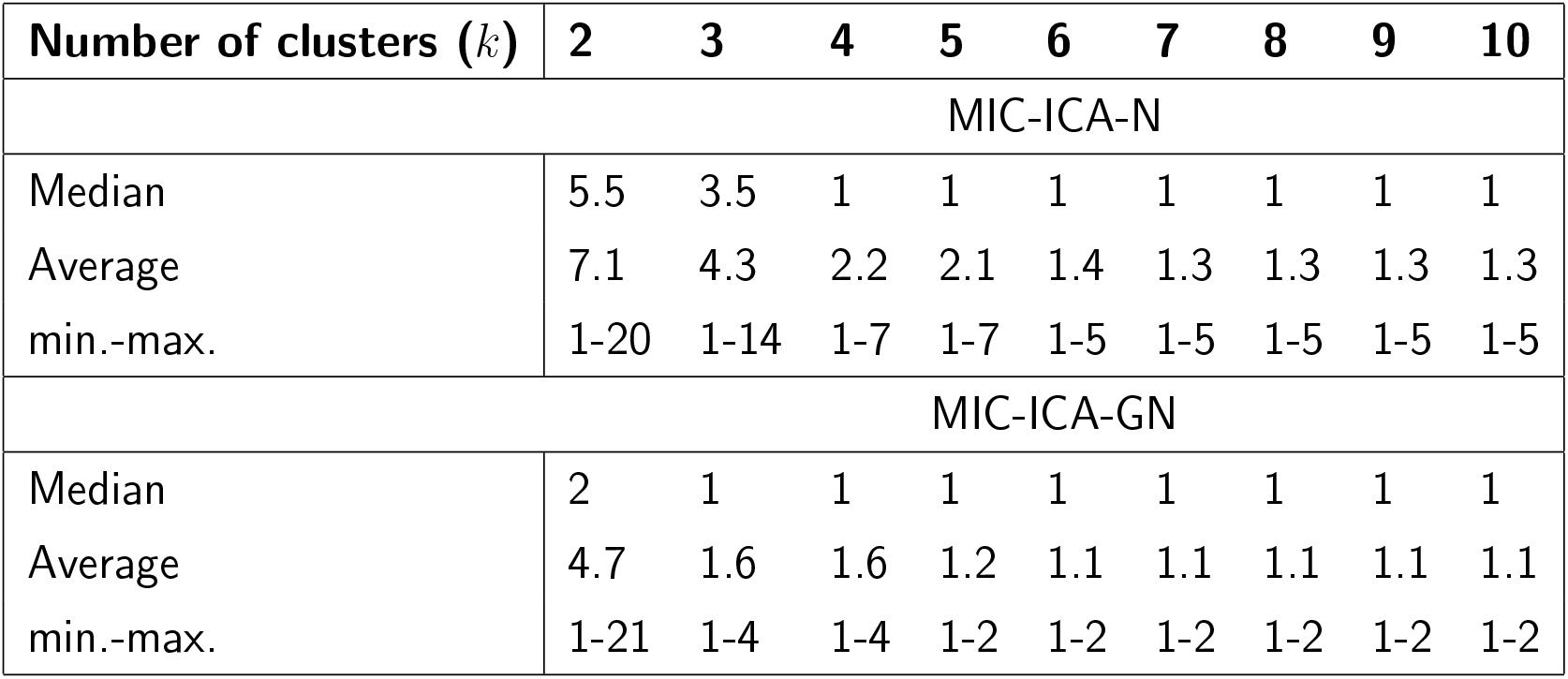
Effect of the number of clusters (*k*) on the number of artefactually classified ICA components.

| Number of clusters ($k$) | 2 | 3 | 4 | 5 | 6 | 7 | 8 | 9 | 10 |
| --- | --- | --- | --- | --- | --- | --- | --- | --- | --- |
| MIC-ICA-N |  |  |  |  |  |  |  |  |  |
| Median | 5.5 | 3.5 | 1 | 1 | 1 | 1 | 1 | 1 | 1 |
| Average | 7.1 | 4.3 | 2.2 | 2.1 | 1.4 | 1.3 | 1.3 | 1.3 | 1.3 |
| min.-max. | 1-20 | 1-14 | 1-7 | 1-7 | 1-5 | 1-5 | 1-5 | 1-5 | 1-5 |
| MIC-ICA-GN |  |  |  |  |  |  |  |  |  |
| Median | 2 | 1 | 1 | 1 | 1 | 1 | 1 | 1 | 1 |
| Average | 4.7 | 1.6 | 1.6 | 1.2 | 1.1 | 1.1 | 1.1 | 1.1 | 1.1 |
| min.-max. | 1-21 | 1-4 | 1-4 | 1-2 | 1-2 | 1-2 | 1-2 | 1-2 | 1-2 |

An example of individual source-level activations during vocal and silent naming is depicted in Figure 8. In this subject, the different methods removed 1, 1, and 2 components (MIC-ICA-N, MIC-ICA-GN, Manual-ICA). For the individual, the source-level activation patterns did not differ much between the different cleaning methods, while the original and cleaned data always differed clearly from each other. In contrast, the cleaning did not change the activation patterns for the silent naming data.

**Figure 8.**
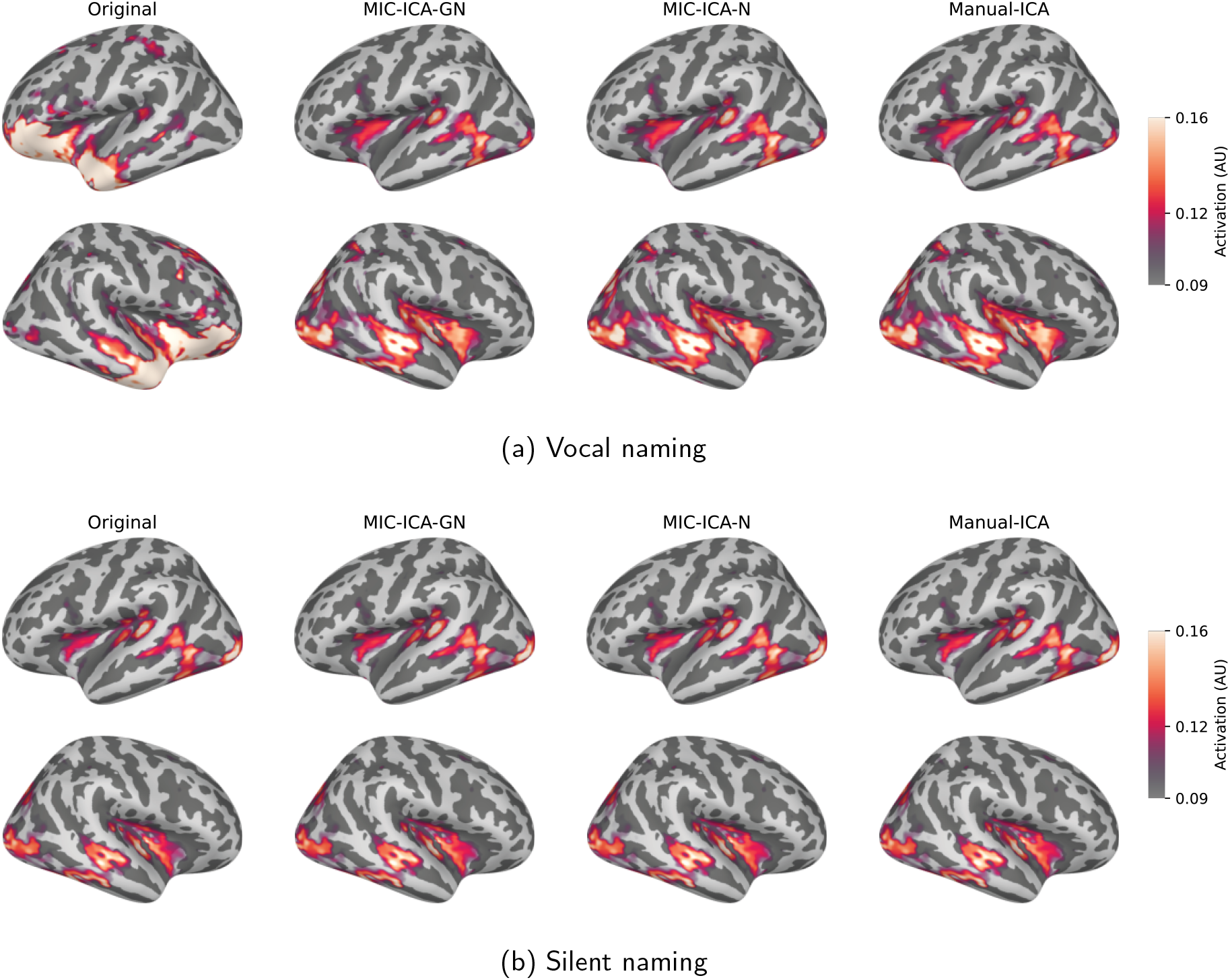
Source-level results in one individual during the vocal and silent naming tasks. The leftmost column shows the original responses and the three other columns show the cleaned activity after the three different artefact removal methods.

### 3.2 Comparison between different artefact removal methods

#### 3.2.1 Sensor-level results

Figure 9 shows the sensor-level data averaged over all subjects during the vocal and silent naming tasks (0-1500 ms) before and after the cleaning procedures. In the vocal naming task, most of the gradiometer elements (either one or both of the sensors in the pair) differed statistically (60/61/60 elements for MIC-ICA-GN/MIC-ICA-N/Manual-ICA of which 13/11/14 were pairs) between the original and the cleaned data (*p <* 0.001). Significant differences between the original and cleaned evoked responses were found in the time intervals of 140-1230 ms, 140-1240 ms, and 140-1240 ms for MIC-ICA-GN, MIC-ICA-N and Manual-ICA, respectively. For the silent naming task, there were no statistically significant differences between the cleaned and original evoked responses for any of the methods.

**Figure 9.**
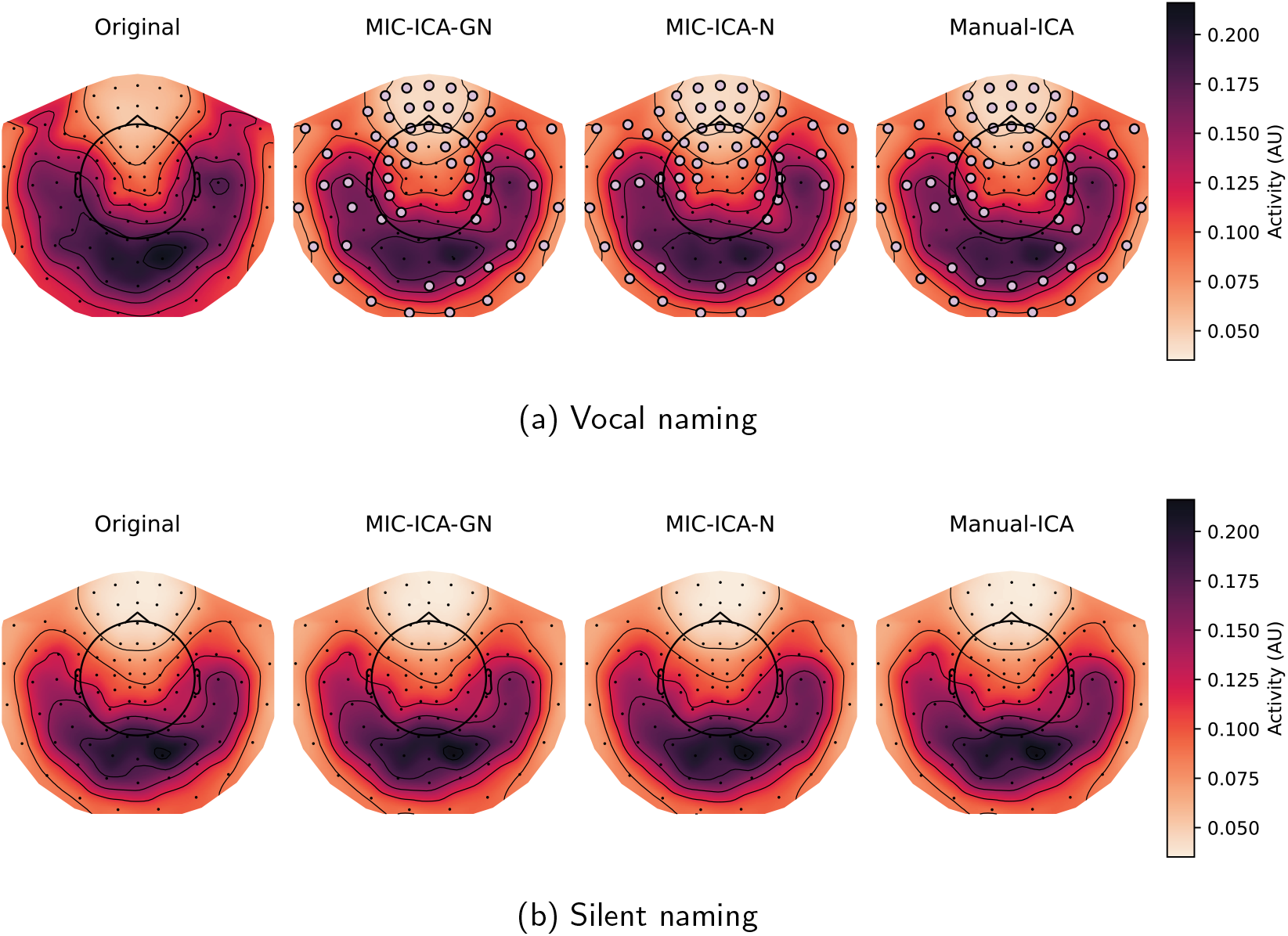
Sensor-level topographies of the vocal and silent naming data before and after cleaning. The data is averaged over all subjects in the time window of 0-1500 ms. The leftmost column shows the original response and the other columns show the cleaned activity during the vocal or silent naming tasks for the three different artefact removal methods. The sensor elements marked with dots mark the statistically significant differences between the cleaned vs. original data.

To describe the removed signal in more detail, *RMSD*_*ch*_ values comparing the original and the cleaned evoked responses in the vocal naming task were calculated for each subject independently and averaged into a group-level topography map (Figure 10). The obtained topography maps, describing the amount of removed artefact, were highly similar across methods. The *RMSD*_*ch*_ topographical maps have 5 local maxima (see Figure 10). Two of these maxima, one on the left side and the other on the right reside over areas that are considered to be relevant for language processing. The evoked responses at these sensor pairs are presented in the inset below and above the topographic maps in Figure 10. The grey shadowed area in the insets marks the statistical difference (*p <* 0.001) for the cleaned and original evoked responses of vocal naming task. The cleaned evoked response of each method seems to be similarly different to the original evoked response. Additionally, particularly around 500 ms after the stimulus onset, the cleaned vocal naming evoked responses follow closely the original silent naming responses.

**Figure 10.**
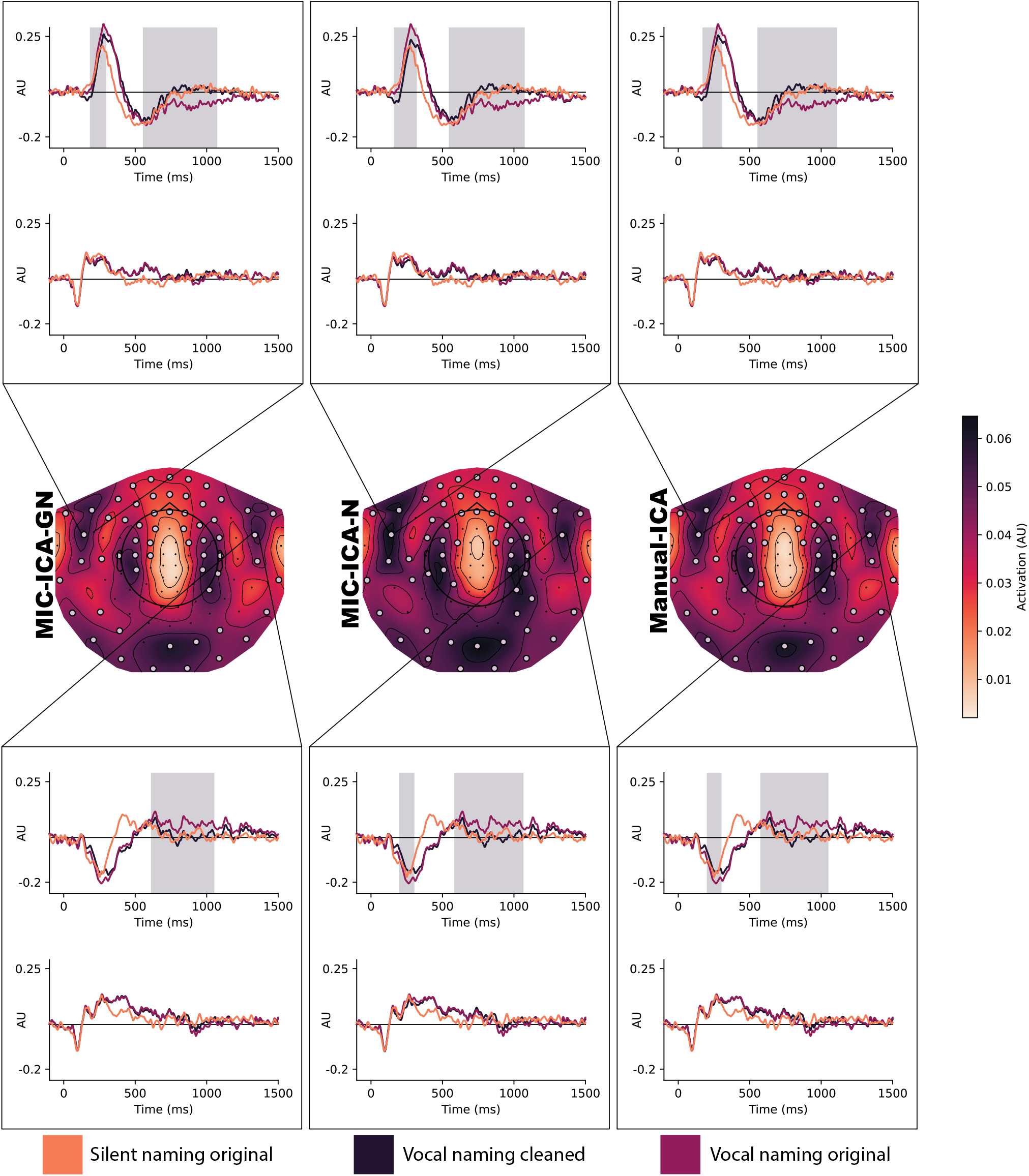
RMSD topography of the removed artefact. The middle row shows the *RMSD*_*ch*_ values at each MEG sensor pair illustrating the removed amount of artefact from the vocal naming data. Sensors that had statistical differences between the original and cleaned data are highlighted in the topographies. The insets present the evoked responses at two sensor pairs residing over areas that are relevant for language processing. The grey area in the insets shows the time during which the original and cleaned vocal conditions differed significantly.

## 4 Discussion

Studying speech production with neuroimaging methods is complicated by the presence of significant artefacts in the measured brain activity during vocal speech. This study aimed to develop an ICA-based speech artefact removal routine for MEG data by a) using EMG data measured from facial muscles during a facial gesture task for isolating the speech-induced artefacts, and b) automating the artefact component selection by utilizing mutual information (MI) and independent component (IC) approaches. Our results show that the automated MIC-ICA approaches produced highly similar results to the manual selection of artefact components. Additionally, our findings indicate that the developed facial gesture task increased the reliability of the automation. The source-level results demonstrate that the artefact removal procedure significantly changed the brain activation, especially over the anterior temporal lobe.

Previous studies have addressed the problem of speech artefacts by modifying the required speech response, for example by using delayed (Ala-Salomäki et al., 2021; Jescheniak et al., 2002) or silent naming (Eulitz et al., 2000; Liljeström et al., 2009). Compared to these approaches, the present work used an immediate vocal response, which has significant benefits. During a vocal task, it is, e.g., possible to monitor that subjects follow the task instructions correctly and identify potential speech errors. Similarly to previous solutions (Alexandrou et al., 2017; Barbati et al., 2004; Henderson et al., 2013; Porcaro et al., 2015; Vos et al., 2010), artefact removal was here based on ICA. However, the automated component selection in this study used MI between the EMG and IC signals. The use of MI was motivated by its capability to measure both linear and non-linear relationships in addition to its potential to detect high-frequency artefacts better. MI can be especially beneficial if the EMG measures are not comprehensive of all the muscle activations contributing to the artefacts, as correlation may fail in this respect. As speech artefacts are known to arise from many different sources, measuring speech artefacts more comprehensively may not be feasible: using only a few surface EMG electrodes may not be enough to address even the majority of speech artefact sources, and adding more EMGs is typically impractical.

Overall, the MIC-ICA-GN and MIC-ICA-N seemed to produce similar results to the Manual-ICA, emphasising their benefits: reducing subjective observer bias and speeding up the artefact removal process. Neither of the automated methods clearly outperformed the other, but the MIC-ICA-GN had several advantages. The MIC-ICA-GN method affected the least the activation in the areas typically considered important to speech production, e.g., the ventral frontal lobe (Levelt et al., 1998). In addition, MIC-ICA-GN affected the left and right hemispheres more symmetrically, suggesting that the ICA decomposition could better find the speech artefact with the additional facial gesture task. MIC-ICA-GN was also found to be less sensitive, in respect of the number of removed components, to the selected number of clusters *k*, which in turn suggests that the MIC-ICA-GN is more robust than the MIC-ICA-N. Since MIC-ICA-N could be considered to fail for a few subjects, there is uncertainty in using this method. Therefore, this work promotes the use of a gesture task as a part of the study protocol, which enables reliable automation for speech artefact removal. However, the MIC-ICA-N method can still be used to speed up the removal of speech artefacts, for example in combination with visual inspection of components. Despite these small differences, it should be emphasized that there were no visible differences in the group-level brain activation or statistical differences between the methods in pair-wise testing.

In the sensor-level analysis, the artefact reduction affected the majority of the sensors regardless of the artefact removal method. These procedures, however, did not affect the MEG signals earlier than around 140 ms, suggesting that no significant signal was removed in the early time windows. In addition, the artefact-free vocal naming and silent naming responses were largely similar to each other, especially after 500 ms. Before 500 ms some differences were detectable, but they might also relate to the differences in the brain processes underlying vocal and silent naming. It can be expected that the brain activation during vocal naming involves, for example, preparation for articulation and motor planning in addition to the vocal output (Levelt et al., 1999).

A limitation of the current study lies in the fact that all chosen methods use ICA, and there is no certainty that the ICA separates sources perfectly, such that no brain activation is mixed with the artefactual components. Furthermore, the true brain activation during speech is not known, introducing uncertainty in the evaluation of the results. Nevertheless, Manual-ICA can be taken as the current golden standard for cleaning the speech artefacts without addressing how perfectly it separates the artefact and the speech-related brain activations. Understanding this limitation, our study does not claim that MIC-ICA-GN perfectly cleans the measured MEG data from speech artefacts, but it allows to obtain results that are as good as the results from Manual-ICA in an automated way. This conclusion is supported by the results demonstrating that the artefact-free vocal naming data closely resembled the silent naming data.

The EMG sensor placements used in this study have been previously used in M/EEG speech artefact studies (Abbasi et al., 2021; Porcaro et al., 2015), and in speech recognition studies (Meltzner et al., 2018). However, the optimal EMG placement to measure speech might not be the optimal placement for finding the artefact sources, since EMG signals that recognize speech well do not necessarily indicate high mutual information with the MEG artefact sources. Similarly to the EMG placement optimization, the gestures used here are known to produce strong muscular activation (Stepp, 2012), but whether all of them are important or needed to best imitate the speech artefact and to automate the artefact reduction was not addressed here. Future studies could address the optimal EMG placement and gesture task duration and structure. This study provides the basis for their development, as well as for a reliable evaluation of the results.

Notwithstanding these limitations, this study suggests that using the designed facial gesture task and simultaneous EMG measurements together with language-related MEG tasks improves the quality of the automated speech artefact removal process. Furthermore, the use of mutual information and clustering can help in providing an automated solution for the artefact component selection.

## Author contribution

S.T., S.A., H.R. and M.L. conceived and designed research; S.T. and S.A. collected the data; S.T. and S.C. developed software; S.T. analyzed the data; S.T., S.C., S.A., P.L., H.R. and M.L. interpreted the results; S.T. drafted manuscript; S.T., S.C., S.A., P.L., H.R. and M.L. edited and revised the manuscript.

## Funding and acknowledgements

This research was funded by the Swedish Cultural Foundation (to M.L.), Päivikki and Sakari Sohlberg Foundation (to H.R.), Jenny and Antti Wihuri Foundation (to S.F.C.), State Research Funding to Helsinki University Hospital (grant number TYH2022224 to M.L., P.L., and H.R.) and Research Council of Finland (grant numbers 321460 and 355409 to H.R. and 357660 to P.L.), as well as the Flagship of Advanced Mathematics for Sensing Imaging and Modelling grant 359181.

